# Context-dependent selection as the keystone in the somatic evolution of cancer

**DOI:** 10.1101/542290

**Authors:** Vibishan B., Milind G. Watve

**Affiliations:** Department of Biology, Indian Institute of Science Education and Research (IISER), Pune

**Keywords:** Somatic evolution, mutation accumulation, epidemiology, cancer etiology

## Abstract

Somatic evolution of cancer involves a series of mutations, and attendant changes, in one or more clones of cells. A “bad luck” type model assumes chance accumulation of mutations. The clonal expansion model assumes, on the other hand, that any mutation leading to partial loss of regulation of cell proliferation will give a selective advantage to the mutant. However, a number of experiments show that an intermediate pre-cancer mutant has only a conditional selective advantage. Given that tissue microenvironmental conditions differ across individual organisms, this selective advantage to a mutant could be widely distributed over the population of organisms. We evaluate three models, namely “bad luck”, context-independent, and -dependent selection, in a comparative framework, on their ability to predict patterns in total incidence, age-specific incidence, and their ability to explain Petos paradox. Results show that context dependence is necessary and sufficient to explain observed epidemiological patterns, and that cancer incidence is largely selection-limited, rather than mutation-limited. A wide range of physiological, genetic and behavioural factors influence the tissue micro-environment, and could therefore be the source of this context dependence in somatic evolution of cancer. The identification and targeting of these micro-environmental factors that influence the dynamics of selection offer new possibilities for cancer prevention.

**Highlights:**

- We use a comparative modelling framework to assess the relative importance of random mutations, clonal expansion and context-dependent selection.
- Somatic evolution of cancer is primarily selection-limited, rather than mutation-limited.
- Selective forces based on microenvironmental context vary between individuals. This population-level variance explains most epidemiological patterns.
- Understanding and controlling micro-environmental conditions offers novel avenues of cancer prevention and therapy.

## 1 Introduction

Mathematical models of somatic evolution in cancer have been in development for the past several decades, with a strong focus on mutational processes^1–4^. Most prominently, Tomasetti et al. have argued that cancer risk is largely determined by random mutations^5,6^. On the other hand, the question of somatic evolution in cancer parallels an old debate in the theory of evolution as to how the simple process of random mutations alone can lead to complex structures such as the eye which require the coordinated action of several genes, often perceived as the monkey-on-a-typewriter paradox^7^. The problem of cancer is qualitatively similar to this, but quantitatively even more difficult, since most cancers must evolve independently in each individual. All cancers are necessarily a combination of different types of genomic changes including point mutations, aneuploidy and other chromosomal aberrations, and the cancer phenotype has a large number of distinguishing characeters, encapsulated by the notion of the “hallmarks” of cancer^8–10^. The wide range of characterisitics that these hallmarks include make it astonishing that so many alterations in cell properties come together in cancers purely out of chance.

Selection on intermediate mutants was the logical solution provided for the monkey-on-a-typewriter paradox, and clonal expansion paralleled such a solution in cancer evolution. Every oncogenic mutation on the way to a cancerous phenotype causes the mutant clone to expand, and as the mutant population increases, the probability of a second oncogenic mutation increases proportionately^11^. Implicit in this theory is the assumption that every oncogenic mutation has a selective advantage over the normal cell. Since most changes involved in carcinogenesis relate to evading growth regulatory mechanisms, it is considered logical that any mutation that allows for such evasion will have a natural selective advantage. However, evidence has been accumulating over the past few years that the fitness advantage of a mutant is largely dependent on the tissue micro-environment^12,13^. Studies in mice^14^ and humans^15^ have demonstrated the effect of contingent factors, such as behavioural profiles and lifestyle parameters, on cancer progression. These findings provide clear indications that the selective forces which determine mutant clone fitness can vary considerably across individuals, leading to *context-dependent clonal expansion* of potentially oncogenic mutants. The role of such differential selective forces in somatic evolution has not been adequately addressed by cancer models so far.

In this paper, we compare three modelling approaches, two of which explicitly include selection at different levels: (1) random mutagenesis, or the “bad luck” hypothesis, (2) context-independent expansion of mutant clones within individuals, and (3) context-dependent selection, where the selective forces vary across individuals. Genetic mutations are a fundamental aspect of multi-stage models of carcinogenesis. In our models however, we use the term mutation more broadly, to denote any change that is heritable within a given cell lineage, genetic or epigenetic.

Since well-curated data are available for human cancer incidence patterns through the SEER databases^16^, we develop models of these three processes, and compare their predictions with the epidemiological picture of cancer in the human population. A good working model of somatic evolution of cancer needs to explain the following epidemiological features:

1. Total and age-specific incidence: Total incidence of cancer across types lies below 30%, while age-specific and cumulative incidence patterns show more variations between cancer types^16^. Interestingly, recent analyses have shown for several cancers that the age-specific incidence rates decline late in life^17^, in contrast with general model predictions of a power law increase in incidence with age. Some theoretical explanations for this decline have been suggested^18^, but exact causes remain unclear. The late-life decline causes the cumulative incidence to saturate with age at a small percentage of the population size. No matter the lifespan, the proportion of cancer in the population can never reach 100%, which represents a finite limit that is not determined by time.
2. Incidence vs cell number: The relationship between cancer risk and cell number (as the lifetime number of cell divisions, *lscd*) has been kept in the spotlight by recent work by Tomasetti et al.^5,6^. Although the linearity of the relationship between *lscd* and cancer risk is still under debate, available data can be used to comparatively test model predictions.
3. Incidence vs mutation rate: Empirical data on this relationship are less common, as mutation rates are difficult to measure reliably, although some efforts have been made^19^. However, a general notion exists that higher mutation rates increases cancer risk, and this remains to be tested rigorously, barring theoretical work dealing with the effect of mutagens on patterns in incidence data^18^.
4. Non-mutagenic carcinogens: There are several agents, including hormones and growth factors, that increase cancer risk without affecting the basal mutation rate^20^. The activity of these agents, and their signature in epidemiological patterns, are both important in building a complete framework of explaining cancer etiology.
5. Peto’s paradox, and similar observations: This relates to the incidence-cell number relationship, as cancer risk is seen not to scale with body size or cell number across species^21^, but may correlate with the latter within a species^22^. A wide range of explanations have been offered for this observation^23^, and in a modeling context, these explanations can come from the model itself (intrinsic), or involve extrinsic factors, such as evolved cancer defences, that are not part of the model framework.

## 2 The “bad luck” model

This hypothesis assumes that the required set of driver mutations accumulate in a cell by chance alone. This may happen over a period of time, or in a single large-scale event, like chromothripsis^24^. Regardless of whether mutations accumulate sequentially or otherwise, the “bad luck” model does not assume selection of any kind on mutations over the course of somatic evolution.

Consider an organism with a population of *n* stem cells, each with a mutation rate per cell generation per locus, *p*. The probability that at least one cell acquires one mutation at a given point of time can be given as 1 – (1 – *p*)^*n*^. If *k* such mutations are requried for cancer onset, the probability of cancer according to the bad luck model can be given similar to (ref. [25]):

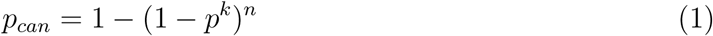

Given the probability of cancer per unit time, *p_can_* from equation 1, the cumulative incidence of cancer for age, *A*, can be expressed as below:

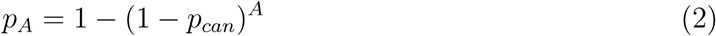

From equation 1, it is clear that the probability of cancer has a threshold relationship with both *n* and *p*, such that incidence rises from near zero to 100% over a small range of *p* and *n*, as shown in Figure 1.

**Figure 1:**
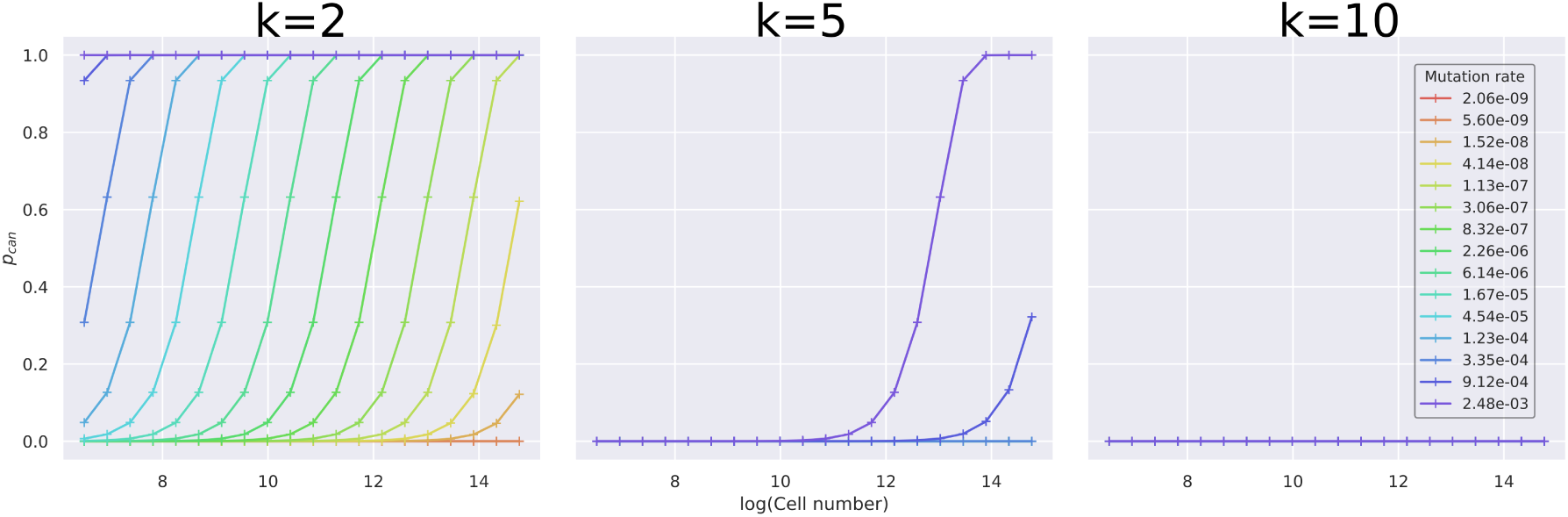
Cancer probability, *p_can_* vs cell number for the “bad luck” model; *p_can_* remains near zero for small *n*, and rises to one over a narrow region of *n*. This proability is cumulative and therefore reflects total incidence in the population. The number of oncogenic mutations required for cancer onset (*k*) does not affect the existence of a threshold with *n* and *p*, but does affect where the threshold occurs in the parameter space; for *k* = 10, the threshold does not occur within the tested range of *n* and *p*. The legend shows values of *p* for each curve.

From equation 2, the relationship of *p_can_* with age is a monotonically increasing function with saturates only at unity. Figure 2 shows this relationship across the entire parameter range of *n* and *p*, for which this prediction holds. Cancer probability increases monotonically with age, and only saturates at 100% incidence, which stands in stark contrast to the observed late-life decline in age-specific rates. Given a finite lifespan, cancer incidence lies in a realistic range only over a narrow range of *p* and *n*.

**Figure 2:**
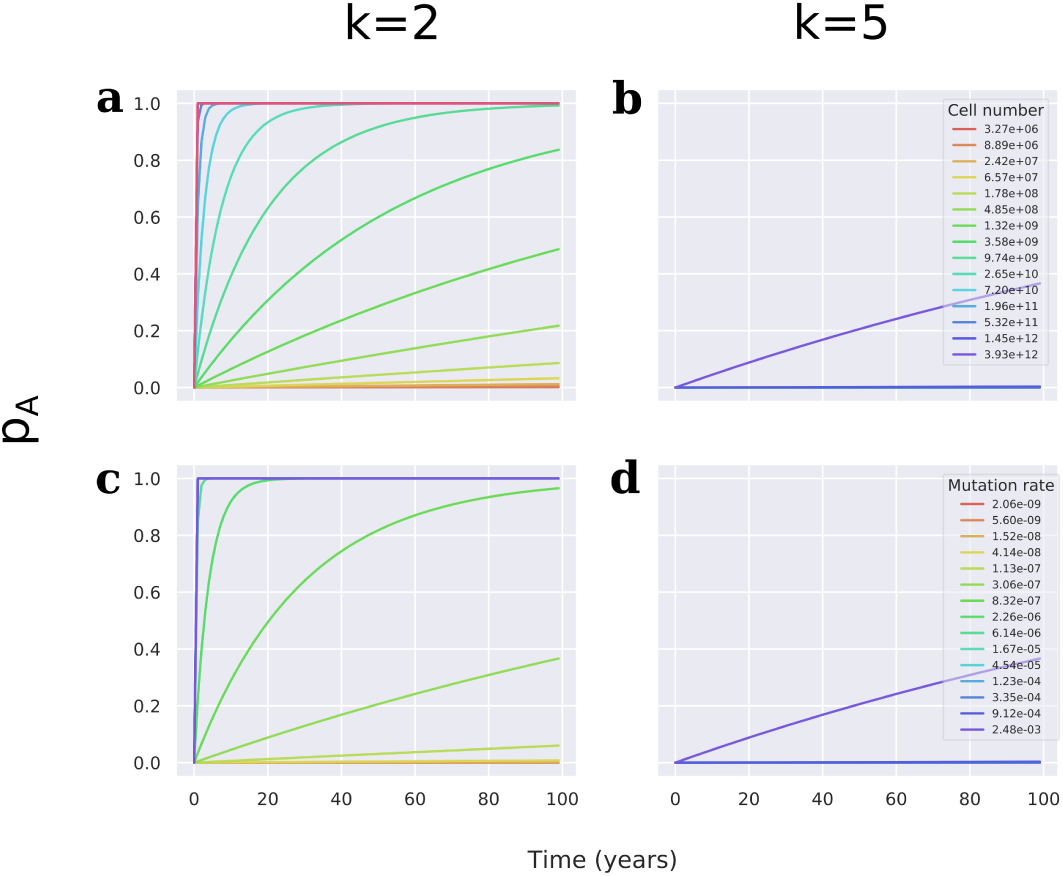
Cancer probability vs age, as given by equation 2, (**a** and **b**) over the range of *n*, and (**c** and **d**) over the range of *p*; inset legends give the corresponding values of *n* and *p*. Cancer probability increases monotonically with age, saturating only at one in most cases. Where the probability does approach realistic values of total incidence, it still does not reflect the late-life decline in incidence rates observed epidemiologically. As in Figure 1, the number of oncogenic mutations required for cancer onset does not change the nature of the incidence-age relationship. For **a** and **c**, *k* = 2, and for **b** and **d**, *k* = 5.

Including the cost of lethal/deleterious passenger mutations does not qualitatively alter the general prediction of thresholds with *n* and *p*, although the position of the threshold may change, as has been observed independently^2^. Taken together, this formulation of the “bad luck” model predicts a sharp threshold relationship of cancer probability with both *n* and *p*, as opposed to the observed progressive increase over several orders of magnitude^5,6^. For values of *n* and *p* around this threshold, cancer probability increases rapidly with age but is inconsistent with late-life decline in cancer incidence.

## 3 Models with selection on mutants

For models that involve selection, we choose to explore the effects of context-independent clonal expansion and context-dependent selection through a simulation-based stochastic framework.

We use a linear process to model the sequential accumulation of mutations in a population of stem cells. We begin by considering the development of a generalized tissue compartment in each organism starting from one stem cell, with mutation rate per cell generation per locus, *p*, growing logistically to a carrying capacity, *n*, following the discrete logistic equation below:

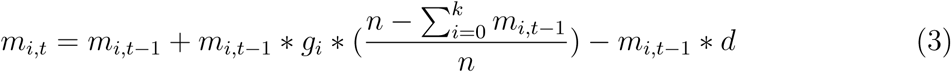

Here, *m_i,t_* is the size of the *i*th mutant population at time, *t*, with *i* = 0 being the non-mutant cell population, *g_i_* is the corresponding logistic growth rate, *d* is the common death rate, and *k* is the threshold number of oncogenic mutations required for cancer onset. As the organism develops into an adult, net growth in the stem cell compartment saturates, but reaches a dynamic equilibrium between cell death and renewal. The stem cell population can be reduced, either by death of stem cells or differentiation, as reflected by *d* in equation 3. Assuming a common death rate for all cell populations, the replacement of the lost cells by either mutants or non-mutants is a function of their growth rates. We simulate new mutation events stochastically; the probability of at least one cell mutating is given by 1 – (1 – *p*)^*m_*i,t*_*^, and if this probability exceeds a random number between 0 and 1, a new (*i* + 1)th mutant population is initiated. Each new oncogenic mutation could give a growth advantage over older cell populations, leading to successive cycles of clonal expansion in which the newer population gradually replaces older cells through competitive exclusion. We simulate this linear evolution process until *k* mutations have been accumulated, which is the assumed threshold for cancer onset. Death of the individual occurs either at cancer onset when the *k*th mutation occurs, or at the end of the assumed lifespan of 100 years, whichever happens first. This simulation is repeated independently for a population of 10000 individuals, and the population-level cancer incidence is recorded, along with the age of onset.

### 3.1 Choice of parameter range

In order to standardize the discrete logistic simulation, we assume the time unit to be one day per logistic growth step. Most human organs complete development and maturation wihtin the first 10-20 years of the lifespan, and the final carrying capacity achieved is the adult stem cell number, ranging between 10^6^ and 10^11^ across different tissues (supplementary material from^5^). Given the final population size and the time taken to reach it, a simple calculation based on the logistic equation shows the required growth rate for a non-mutant stem cell to be in the range of 0.00383-0.0131. Starting from the non-mutant growth rate, *g*_0_, growth rates are assumed to increase linearly for each subsequent mutant population, with the slope given by 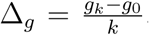. Ranges of *n* and *p* are retained as in the “bad luck” model.

### 3.2 The context-independent selection case

While the clonal expansion theory introduced the notion of selective advantages to oncogenic mutants, it makes the implicit assumption that identical mutations have the same selective advantage in every individual in which they occur; stated otherwise, individuals do not differ in their propensity for mutant clonal expansion. To capture this in the context-independent selection case, we use the same slope, ∆_*g*_ for all individuals in the simulation.

### 3.3 The context-dependent selection case

As argued earlier, it is becoming increasingly clear that the competitive outcomes of identical mutations can depend strongly on the micro-environmental context in which cell competition occurs. In order for selection on mutants to be context-dependent in our model, we randomize the slope, ∆_*g*_ over the population from a given normal distribution. Each individual begins with the same *g*_0_, but the progression of growth rates is randomized across individuals, such that individuals with large *g_i_* would progress faster towards cancer onset, while those with small, or negative values of *g*_*i*_ would never progress to a cancerous state as the mutant gets selected against. This produces variation across individuals for cancer propensity.

### 3.4 Predictions from the selection models

As in Figure 3, under the assumption of context-independent selection, the incidence of cancer shows a strong threshold relatioship with age, where up to a certain age, cancer is unlikely or rare, but increases rapidly to 100% within a relatively short span of time. The age-specific incidence declines only when the cumulative incidence approaches 100%, and the age curves show a deterministic sharp threshold despite mutations being stochastic. As with the “bad luck” model, incidence has a threshold relationship wtih both *n* and *p*, as reflected in the maximum cumulative incidence, *I_max_* (Figure 3**d** and **h**). The age at half-maximum incidence is a decreasing function of both *n* and *p* (Figure 3**c** and **g**). Where the incidence of cancer is near the realistic range, for small values of *p* and *n*, the late-life decrease in incidence is still not reflected in the context-independent selection case. The prediction of 100% incidence is due to the fact that all organisms in the population share the same growth rate progression for mutants; giving a distribution to *p* reduces the sharpness of the incidence threshold with age, but saturation of incidence remains at 100% (Figure 4).

**Figure 3:**
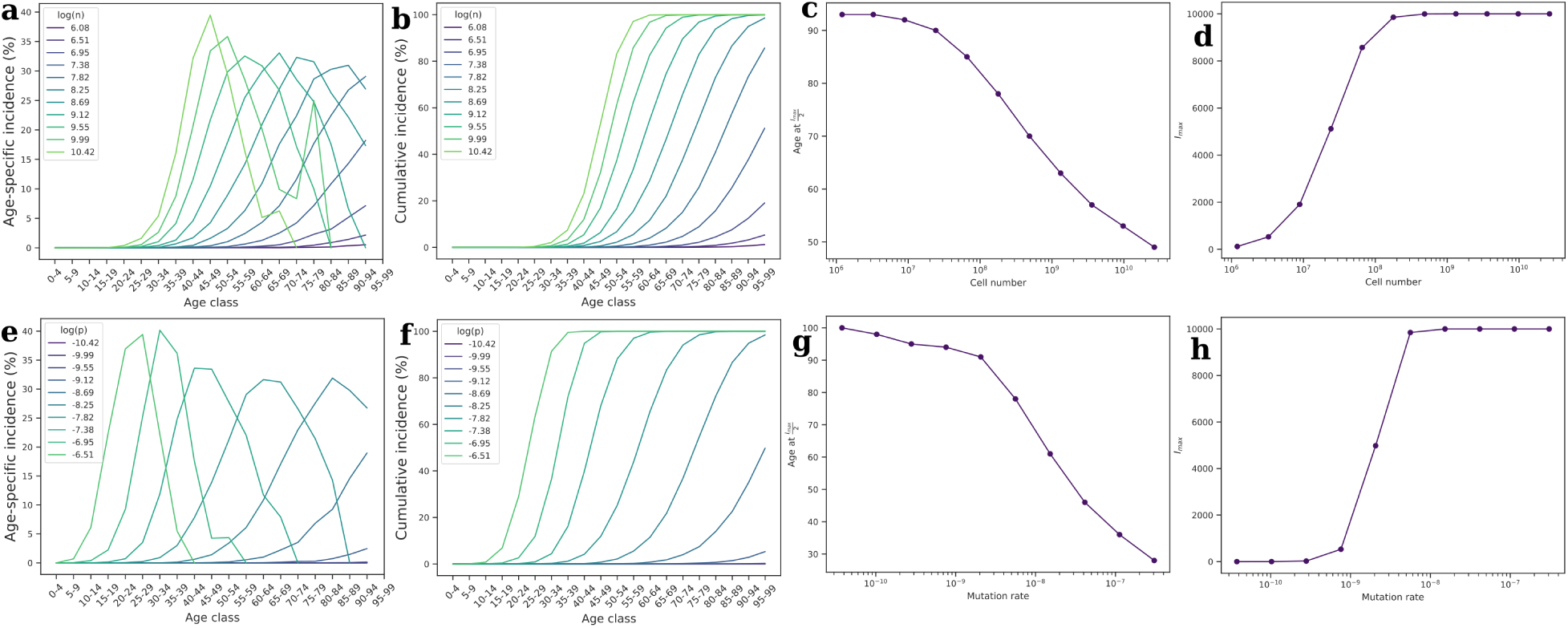
Incidence patterns from the context-independent selection model over the range of (**a**-**d**) *n*, and (**e**-**h**) *p*. From left to right in each row, the plots are of (**a**, **e**) age-specific incidence per 100,000 vs age, (**b**, **f**) cumulative incidence (% of simulated population) vs age, (**c**, **g**) age at which half the maximum incidence is reached vs *n* or *p*, and (**d**, **h**) the maximum cumulative incidence, *I_max_* vs *n* or *p*. Incidence saturates only at 100%, both with time (**b** and **f**), and with *n* and *p* (**d** and **h**). Inset legends for the age curves are *log*(*n*) and *log*(*p*) in the top and bottom row respectively. For **a**-**d**, *p* = 5.603 * 10^*−*9^, and for **e**-**h**, *n* = 1.785 * 10^8^. Growth rates progress linearly in the general form, *g_i_* = 0.007 * (*i* + 1), where *i* = 0*, …, k* and *k* = 5. Here, 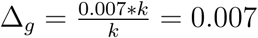.

**Figure 4:**
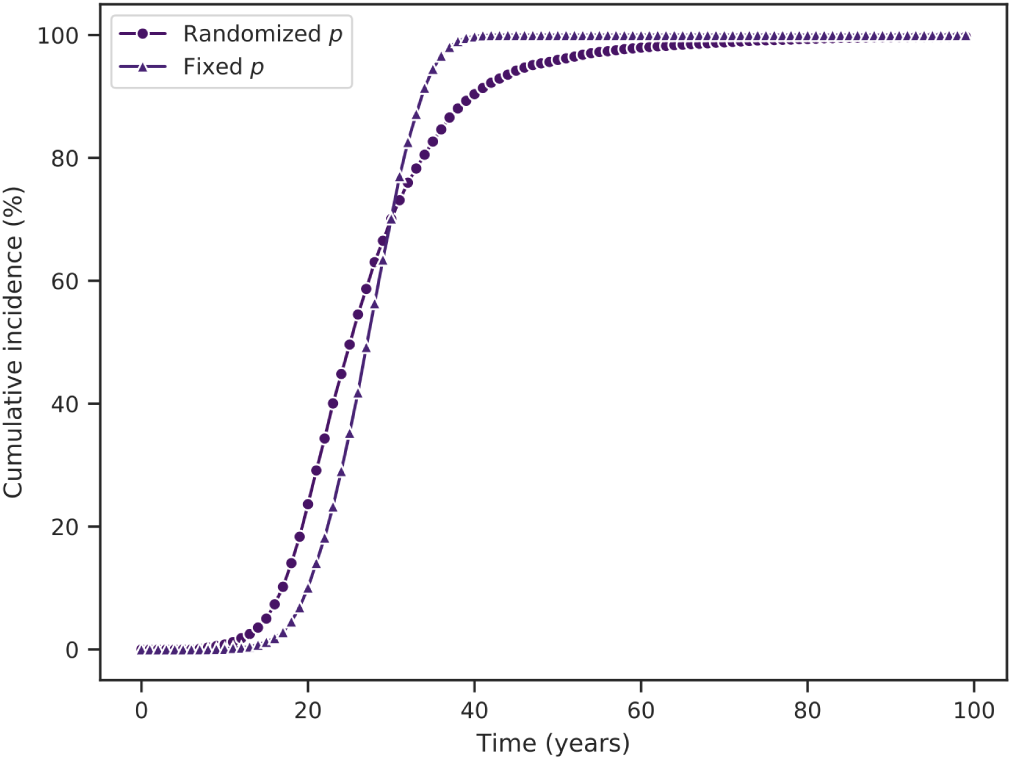
Cumulative incidence (% of simulated population) with randomized vs fixed *p* for the population in the context-independent selection case; giving a distribution to *p* does not reduce the saturation of incidence from 100%, but slows the transition to 100%. In the randomized case, *p* values were drawn from a uniform distribution with range [3.775*10^−11^,3.006*^-7^] and mean, 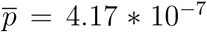, while for the fixed case, *p* = 3.06*10^−7^; for both, *n* = 1.785*10^8^ and *k* = 5.

As opposed to the context-independent selection model, the context-dependent model produces a saturating trend in cumulative incidence that begins to saturate at a level much lower than 100% (Figure 5). This is an important feature of the context-dependent model, as it allows the model to generate more realistic patterns in age-specific cancer incidence. The saturation occurs because propensity for clonal expansion varies across individuals; cancer progression occurs very quickly in some individuals, and not at all in others. Of the three models analysed so far, only the context-dependent model captures a trend similar to the late-life decline observed in many cancers in humans, as the crude incidence curves in Figure 5**a** and **e** show. It is also important to note that cancer incidence increases progressively over several orders of magnitude of both *n* and *p*, as opposed to the thresholds observed in earlier models (Figure 5**c** and **g**).

**Figure 5:**
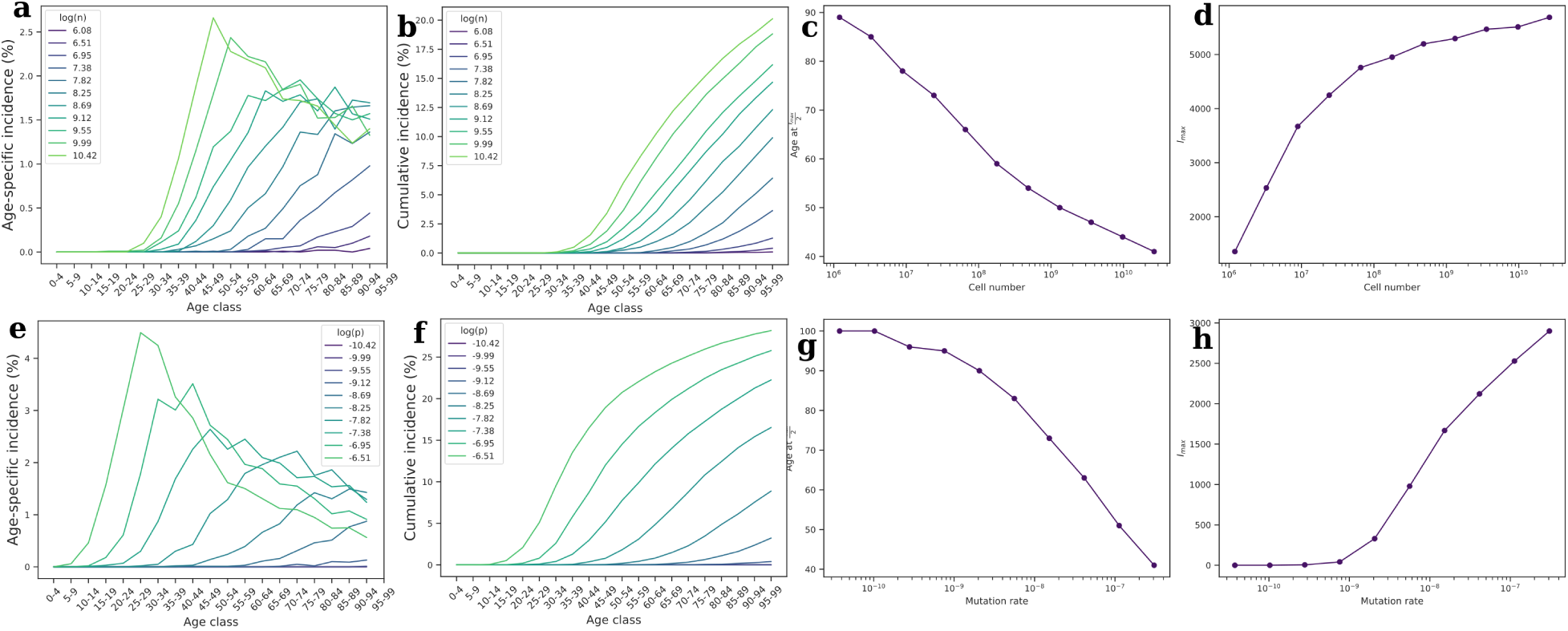
Incidence patterns from the context-dependent selection model over the range of (**a**-**d**) *n*, and (**e**-**h**) *p*. From left to right in each row, the plots are of (**a**, **e**) age-specific incidence per 100000 vs age, (**b**, **f**) cumulative incidence (% of simulated population) vs age, (**c**, **g**) age at which half the maximum incidence is reached vs *log*(*n*) or *log*(*p*), and (**d**, **h**) the maximum cumulative incidence, *I_max_* vs *n* or *p*. As opposed to Figure 3, cumulative incidence saturates much below 100%, both with time (**b** and **f**), and with *n* (**d**), when *g* is randomized in the population. Note that randomization of *p* alone in the population does not reproduce these patterns (Figure 4). Inset legends for the age curves are *log*(*n*) and *log*(*p*) in the top and bottom row respectively. For **a**-**d**, *p* = 5.603 * 10^*−*9^; for **c**-**h**, *n* = 1.785 * 10^8^; and for all figures, *k* = 5. Insets for the incidence curves show *log*(*n*) and *log*(*p*) in the top and bottom rows respectively. Growth rates progress linearly from *g*_0_ = 0.007 to *g_k_* = 0.007 * *µ*, where *µ* is normally-distributed random variable with 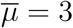 and *σ* = 5. Here, 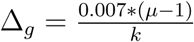.

### 4 Sensitivity of predictions

We relax the quantitative and qualitative assumptions of the model to test whether or not the results are artifactual, contributed by any specific assumption.

Randomizing *k* in the range, [2, 10]^26^, the nature of the relationships did not change qualitatively, although *k* affected the age of cancer onset (Supplementary figures, S2.1-S2.4). As different types of cancers may have different *k*^27^, the effects of *n* and age on incidence trends could also change correspondingly. This is also relevant to the study of similar patterns that do not show up at the whole population level, but may become apparent if the population is stratified by the number of driver mutations, which in turn is known to vary across cancer types.

To test whether the assumption of a normal distribution for ∆_*g*_ is critical for the results, we reevaluated the predictions of the context-dependent selection model for a Gumbel- and uniformly-distributed ∆_*g*_ progression. We find that the shape of the distribution does not affect the saturation of cumulative incidence, or the late-life decline in age-specific incidence predicted by the model, although the shape of the age curve changes (Supplementary material, S1 Figures). We assume generally that ∆_*g*_ is constant for a given individual, but relaxing this assumption by giving a distribution to ∆_*g*_ within each individual would not change the results qualitatively. In such a case, the lowest ∆_*g*_ within an individual would be the rate-limiting step.

Both our selection models assume a linear somatic evolution process, by which the tissue has only one evolving lineage of cells at a time. Alternative modalities have been suggested for tumour evolution, like branched, neutral and punctuated modes^28^, and some modelling effort has gone into studying branched processes in somatic evolution^29^. We do not explicitly model branched somatic evolution in our framework as it is computation-intensive, but our framework is, in principle, with branched and poly-clonal progression. Qualitatively, we do not expect a branched evolution model to alter our primary conclusion that somatic evolution of cancer is largely selection-limited and not mutation-limited. In fact, a branched evolution process allows for a wider range of ecological interactions between sub-clonal populations, mutualistic or otherwise^30,31^. Such interactions again lend into a selection-driven framework rather than a “bad luck”-driven one.

Somatic cell populations are potentially susceptible to mutational meltdown, and a large number of somatic mutations are in fact known to be deleterious. Earlier analysis of the effect of passenger mutations has shown that accounting for them in somatic evolution delays overall incidence of cancer in the population^2^. However, from our context-independent selection case, we see that when total incidence in the population decreases, most cases of cancer are concentrated later in life, similar to the delay in incidence noted before^2^. This pattern is not compatible with the observed late-life decline in incidence. We expect therefore that mutational meltdown due to deleterious passenger mutations is insufficient to explain saturation due to late-life decline in incidence.

## 5 Discussion

Table 1 compares the prediction profile of the three models discussed so far. On the whole, the better prediction profile of the context-dependent model stems from the distribution of ∆_*g*_ in the population. Several growth factors and hormones are known to increase cancer risk, not the least of which is insulin, without increasing the basal mutation rate or the cell number. The action of such “non-mutagenic carcinogens” is not compatible with a mutation-centric approach to carcinogenesis. We propose instead that non-mutagenic carcinogens could act by altering ∆_*g*_. Late-life incidence in the model comes from a slow growth rate progression across mutations, and our analysis suggests that differential progression of mutant growth rates across individuals is sufficient to produce a gradual decline in late-life cancer incidence. This decline is qualitatively different from the age curves of the context-independent model, where incidence drops abruptly over a narrow age range close to 100% incidence. In the context-dependent selection model, the gradual late-life decline in incidence stems from the left-hand tail of the ∆_*g*_ distribution, where some individuals have zero or negative progression of mutations. It becomes important therefore, to consider the local or evolutionary factors underlying such a temporal pattern of mutation accumulation which could potentially altering selective forces in the form of ∆_*g*_. Likewise, the distribution of growth rate progression over the population could also form part of the explanation for saturation of total cancer incidence in the population across tissue types; the negative part of the ∆_*g*_ distribution precludes some part of the population from ever accumulating enough oncogenic mutations for cancer to occur.

**Table 1:** Comparative explanatory power of the three models with respect to observed epidemiological trends

| Epidemiological observation | “Bad luck” | Context-independent selection | Context-dependent selection |
| --- | --- | --- | --- |
| Total incidence across cancer types does not exceed 30% | Saturation at 100% only | Saturation at 100% only | Saturation < 100% possible |
| Age-specific incidence decreases in old age for several cancers | No late-life decline | No late-life decline | Late-life decline predicted |
| Incidence vs $n$ | Sharp threshold saturating at 100% | Sharp threshold saturating at 100% | Progressive increase over several orders of magnitude |
| The phenomenon of non-mutagenic carcinogens | No explanation offered | No explanation offered | Act through $\Delta_g$ distribution |
| Peto’s paradox-cancer risk saturates with $n$ | Evolved cancer defences (extrinsic) | Evolved cancer defences (extrinsic) | Cancer incidence not mutation-limited (intrinsic) |

This line of thinking can also be extended to address Peto’s paradox, which is based on the assumption that cancer is mutation-limited. It is not a new argument that Peto’s paradox can be explained by invoking selection^22,23,32^. The results from our model in fact reinforce the notion that somatic evolution of cancer is largely selection-limited, rather than mutation-limited, which explains why cancer risk does not increase in proportion with cell number and/or body size^21^. Moreover, our analysis introduces the additional dimension of population-level variation in cancer propensity as part of the explanation of Peto’s paradox.

This variation in selection could be attributed to the micro-environment in the tumour or pre-cancerous niche, and includes all the factors that determine the selective advantage of oncogenic or pre-cancerous mutants. Cellular interactions with the extra-cellular matrix (ECM) are known to be important in tissue homeostasis^33^ and the growth of cancer populations^13^. The tissue micro-environment is also a key player in the association between obesity and cancer risk, where obese ECM differs from normal ECM both biophysically^34^ and functionally^35^. The immune system also acts as an important mediator in cancer-related processes; a wide range of immune cells including fibroblasts and lymphocytes are known to influence cancer progression under various conditions^12^, and cancer is increasingly seen as an overactive or abnormal wound healing process^9^. *In vitro*, the IGF-II concentration in culture media is found to markedly alter the selective advantage of an IFG-II over-expressing mutant in cell competition^36^. Pre-cancerous cell lines across cancer stages also show different growth properties depending on growth factor concentration in the culture medium^37^. Similarly, levels of estrogen and progesterone are known to play important roles in regulating mammary cell prolferation^38^, secretion of ECM factors and tumour-stroma interactions^39^, and growth factor production^40^. More recently, experiments have been reported in which behaviourally-enriched environments or physical exercise seemed to show cancer-suppressive effects^14,15^ that correlate with levels of particular growth factors, while studies in mice have also shown substantial variation in tumour sizes induced by identical genetic clones across individuals^41^. These observations together suggest that context-dependent factors from the microenvironment, be they biophysical, ECM-derived, GF-derived, hormonal or immunological, could play a limiting role in cancer progression. An alternative research focus therefore emerges, centered on these microenvironmental factors that drive context-dependent cancer progression, whose identification would lead to novel preventive measures, along with potential implications for therapy.

A composite view of cancer etiology requires not only the incorporation of these, and other complexities in models, but also the comparative testing framework that continues to be rare in cancer literature, barring a few efforts^18^. The value of such a framework is immense, as it allows for falsification of factors that make predictions contrary to observed data; this falsification is frequently more robust and informative than an indirect confirmation of potential causal factors. With our analysis, we hope to bring this framework back into the mainstream at a time when the availaibility of large-scale data spanning levels of biological organization is better than ever before, from the population to the various cellular “-omes”. A comparative framework could prove more powerful now, in the context of robust data and computational techniques, and should therefore become a major focus for the cancer modeling effort.

## Acknowledgements

The authors acknowledge critical input faculty members at the Department of Biology, IISER Pune. We thank Ketaki Bhagwat, Ruchika Kaul-Ghanekar, Prerna Raina, Anil Gore, and Sharmila Bhapat for useful discussions.

