## Supplementary Information for "Context-dependent selection as the keystone in the somatic evolution of cancer"

February 6, 2019

#### S1 Alternative distributions of $g$

As explained in the main text, we explore the effects of using alternative distributions for $\Delta_g$. Figure S1.1 shows the effect of a Gumbel-dsitributed progression, and Figure S1.2 shows that of a uniformly-distributed progression. We note that the different distributions produce qualitatively similar results as the normal distribution used throughout the main text, although the shape of the incidence curves are slightly different.

#### S2 Sensitivity of model predictions to $k$

Keeping with the sensitivity analysis in the main text, we test the effect of $k$ on the observed relationships of $p$ and $n$ with cancer incidence, and discuss the role of $k$ in greater detail. The time taken to cancer onset is a useful parameter in this context as it describes the temporal dynamics of mutation accumulation, while allowing for some limited inference regarding total incidence in the population. If time taken to cancer is short, population incidence is likely to be large in the given parameter space, and vice versa.

Broadly, we find that cancer incidence is significant only up to maximum $k = 10$. Increasing $k$ also reduces total incidence and shifts the observed time to cancer to later in life, as expected based on mutation accumulation. We also find that the magnitude of $k$ could modulate the strength of the association with $n$, and to a lesser extent, with $p$. Importantly, the effect of either $n$ and $p$ is apparent only when taken for one value of $k$ at a time. As Figure S2.1 shows, when time to cancer onset is pooled across all $k$ values, it appears largely independent of either $n$ and $p$. Doing the same for the context-dependent case, we randomized $g$ as explained earlier, along with either $k$ and $n$, or $k$ and $p$. Remarkably, introducing $g$ as a random variable leads to most of the variance in time to cancer being explained by $g$, and to some extent, $n$. Again, the association between $g$ and time to cancer onset is modulated by $k$, as observed for the association with $n$ and $p$. As with Figure S2.1, the effect of $g$ on time to cancer onset is only apparent when considered one value of $k$ at a time (Figures S2.2 and S2.4).

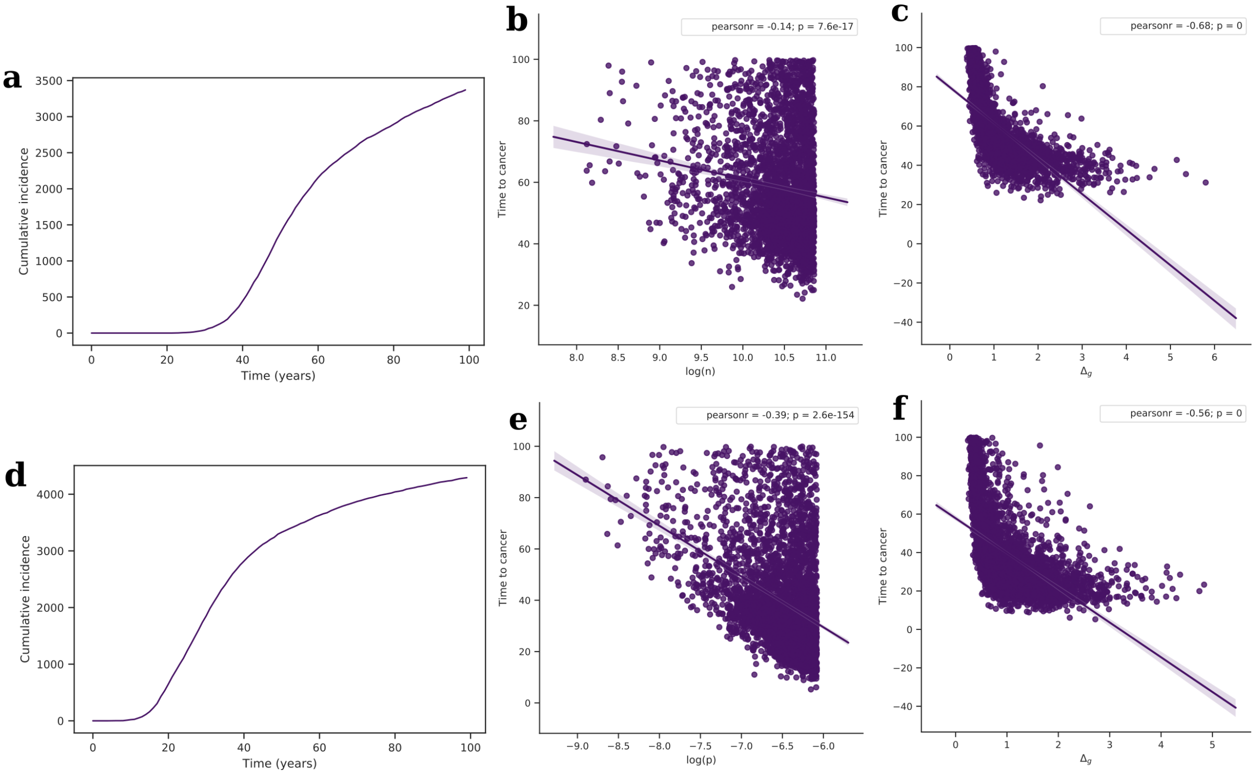


Figure S1.1: $g$ modeled by a Gumbel-distributed random variable, $\mu$, with $\overline{\mu} = 0$ and $\sigma = 3$, co-randomizing $n$ (top row) or $p$ (bottom row), with ranges $[1.203 * 10^6, 2.649 * 10^{10}]$, and $[3.775 * 10^{-11}, 3.059 * 10^{-7}]$ respectively; (**a** and **d**) cumulative incidence (% of simulated population) vs age, and time to cancer vs (**b**) $log(n)$, (**e**) $log(p)$, and (**c** and **f**) $\Delta_g$; $\Delta_g = \frac{g_k - g0}{k}$ and $g_k = 0.007 * \mu$. For all cases, $k = 5$.

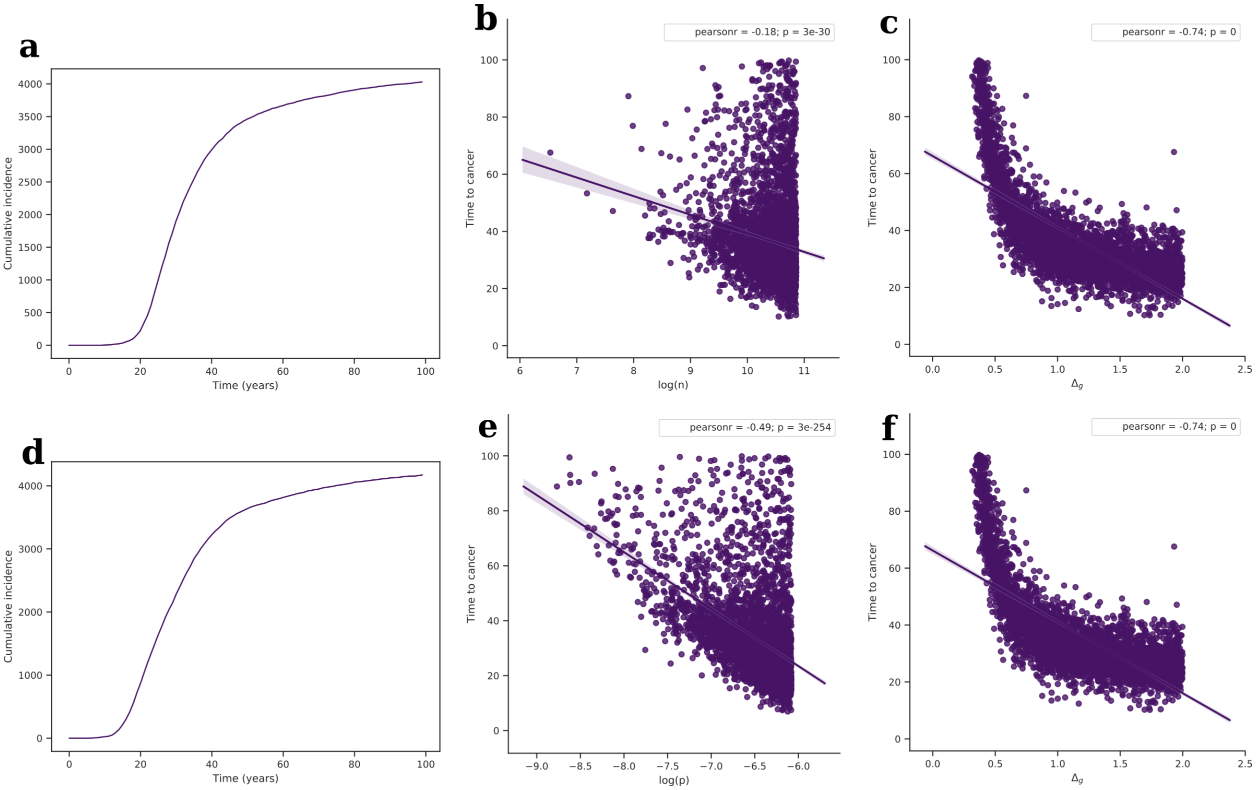


Figure S1.2: $g$ modeled by a uniformly-distributed random variable, $\mu$ with range $[-10, 10]$, co-randomizing $n$ (top row) or $p$ (bottom row), with ranges as specified in Figure S1.1; (**a** and **d**) cumulative incidence for the simulated population vs age, and time to cancer vs (**b**) $log(n)$, (**e**) $log(p)$, and (**c** and **f**) $\Delta_g$; $\Delta_g = \frac{g_k - g_0}{k}$ and $g_k = 0.007 * \mu$. For all cases, $k = 5$.

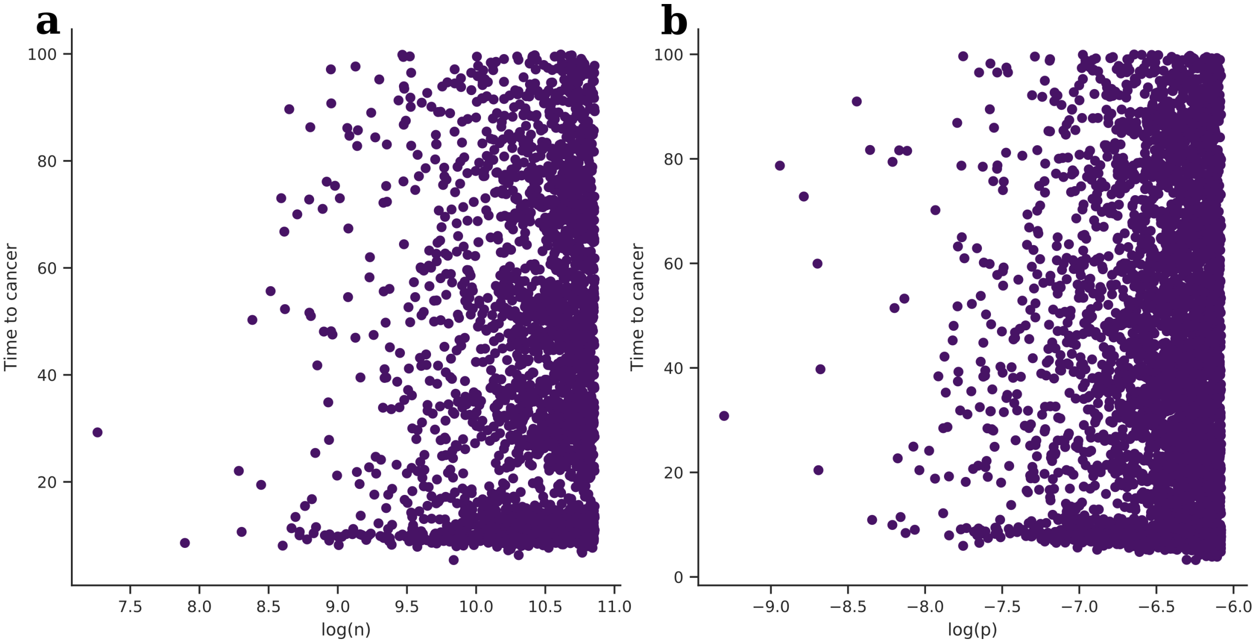


Figure S2.1: Effect of randomizing $k$ in the context-independent selection case. The plots are time to cancer onset against $log(n)$ or $log(p)$, with $k$ randomized with (**a**) $n$, or (**b**) $p$; when time to cancer is pooled across values of $k$, the association between either $n$ or $p$ is practically non-existent. $k$, $n$ and $p$ were uniformly-distributed random variables with ranges $[0, 20]$, $[1.203 * 10^6, 2.649 * 10^{10}]$, and $[3.775 * 10^{-11}, 3.059 * 10^{-7}]$ respectively; for (**a**), $p = 5.603 * 10^{-9}$, and for (**b**), $n = 1.785 * 10^8$.

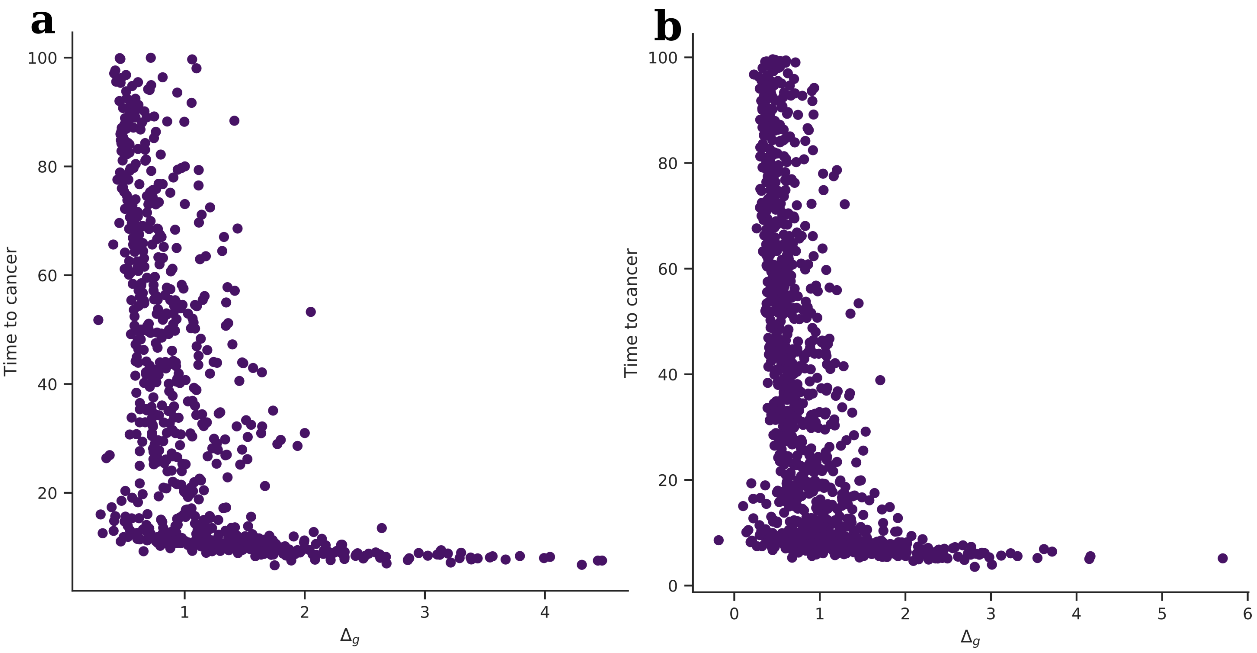


Figure S2.2: Effect of randomizing $k$ in the context-dependent selection case. The plots are time to cancer onset against $\Delta_g = \frac{g_k - g_0}{k}$ and $g_k = 0.007 * \mu$, also randomizing (**a**) $n$, or (**b**) $p$; $\Delta_g$ measures the rate of the growth rate progression in each individual, and $\mu$ is a normally-distributed random variable with $\overline{\mu} = 0$ and $\sigma = 3$. As opposed to $n$ and $p$, the association of $\Delta_g$ with time to cancer is less affected by $k$. When time to cancer is pooled across all $k$, $\Delta_g$'s effect on time to cancer appears distinctly non-linear. $k$, $n$ and $p$ were uniformly-distributed random variables with ranges $[0, 20]$, $[1.203 * 10^6, 2.649 * 10^{10}]$, and $[3.775 * 10^{-11}, 3.059 * 10^{-7}]$ respectively. For (**a**), $p = 5.603 * 10^{-9}$, and for (**b**), $n = 1.785 * 10^8$.

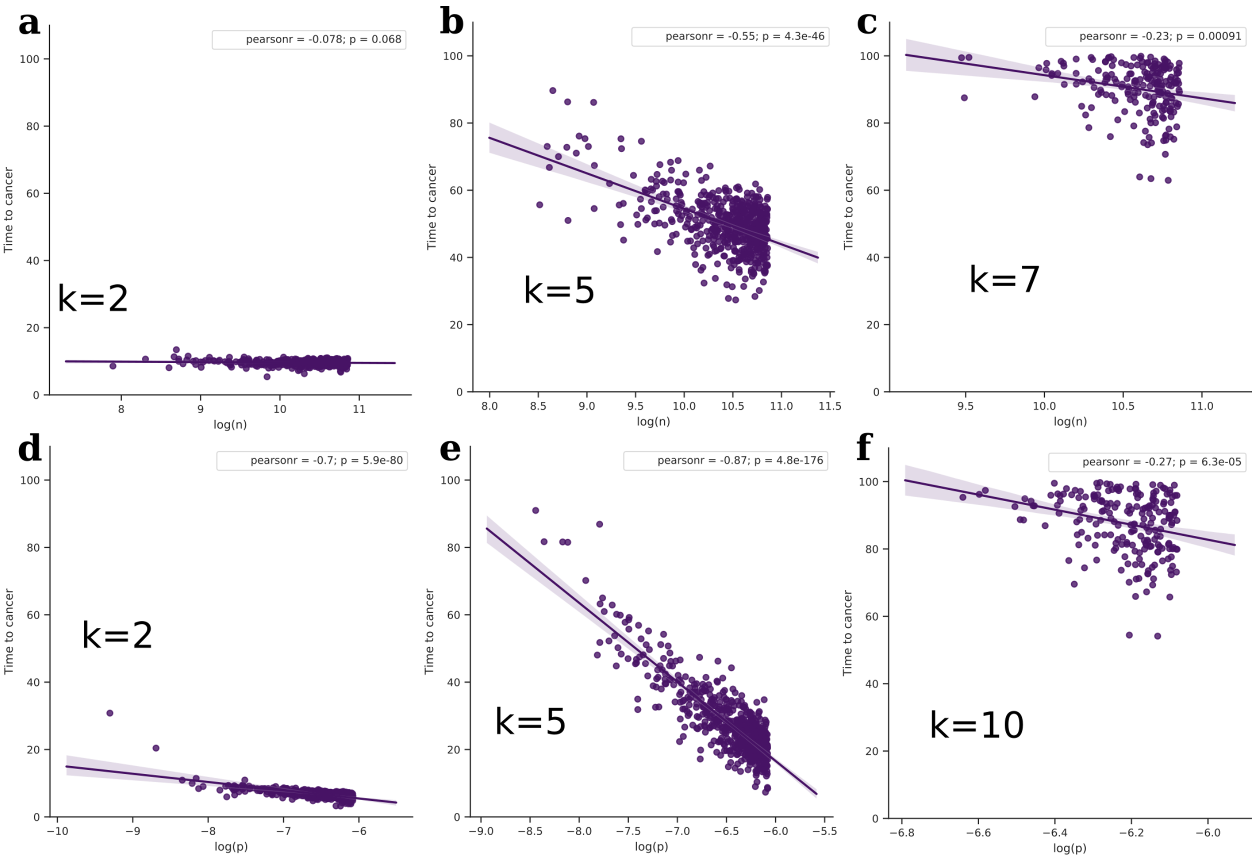


Figure S2.3: Effect of $k$ in the context-independent selection case. The plots are time to cancer onset against $log(n)$ or $log(p)$, with $k$ randomized with (**a**-**c**) $n$, or (**d**-**f**) $p$; value of $k$ in the inset corresponds to the number of threshold oncogenic mutations assumed for the corresponding points. From (**a**-**c**), for higher threshold of oncogenic mutations, the effect of $n$ on time to cancer gets stronger, as shown by the improvement in the association. For small $k$ however, $n$ does not affect the age of cancer onset. On the other hand, $p$ has a strong effect on the time to cancer at every value of $k$ considered. $k$, $n$ and $p$ were uniformly-distributed random variables with ranges $[0, 20]$, $[1.203 * 10^6, 2.649 * 10^{10}]$, and $[3.775 * 10^{-11}, 3.059 * 10^{-7}]$ respectively. For (**a**-**c**), $p = 5.603 * 10^{-9}$. For (**d**-**f**), $n = 1.785 * 10^8$.

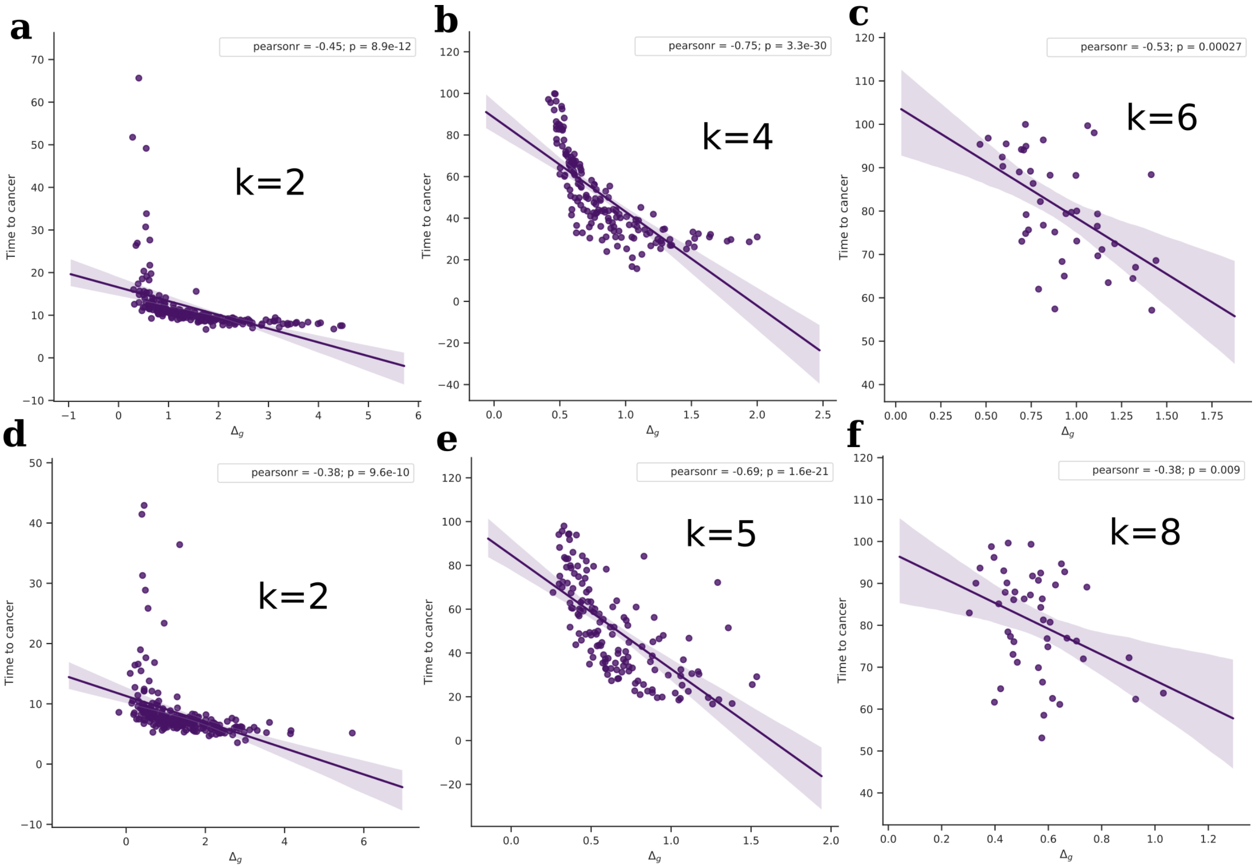


Figure S2.4: Effect of $k$ in the context-dependent selection case. The plots are time to cancer onset against $\Delta_g = \frac{g_k - g_0}{k}$ as defined earlier, with $k$ randomized with (**a-c**) $n$, or (**d-f**) $p$; value of $k$ in the inset corresponds to the number of threshold oncogenic mutations assumed for the corresponding points. Compared to Figure S2.3, $\Delta_g$ explains variance in time to cancer much better than either $n$ or $p$. This is true of both (**a-c**) when $n$ and $k$ are also randomized, and (**d-f**) when $p$ and $k$ are also randomized. The effect of $\Delta_g$ is nevertheless modulated by the required $k$, as reflected by the range of $\Delta_g$ for which cancer occurs; the scale of the x-axis across the figure is indicative of this effect. Ranges of $k$, $n$ and $p$, and the underlying distribution of $g$ are the same as in Figure S2.3. For (**a-c**), $p = 5.603 * 10^{-9}$. For (**d-f**), $n = 1.785 * 10^8$.

### S3 The nature of the relationship between cancer incidence and cell number

As mentioned in the main text, the last few years have seen two prominent attempts by Tomasetti et al. to examine the relative contributions of spontaneous mutations, genetic and environmental factors in cancer development[1,2]. In their analyses, they estimate *l*ifetime *s*tem *c*ell *d*ivisions (*lscd*) for 17 tissue types and correlate them with the incidence of cancer using data from the US, and across the world. In the 2017 paper[2], they use the IACR datasets for cancer incidence that consisted of 423 databases corresponding to different countries, of which 347 had incidence data for all 17 cancer types considered here. With these data, they report a strong statistical association (Pearsons r $\approx$ 0.8) between *lscd* and incidence of cancer in a given tissue type. The correlations are on a log-log scale, and on that scale, they are considerably strong. At face value, this is in line with the expected relationship if cancer arises out of largely random processes of mutagnesis, and the authors therefore view this association as clear evidence for the causal significance of spontaneous mutations in carcinogenesis. They use this association to further attribute the majority of cancer incidence to random replication errors alone.

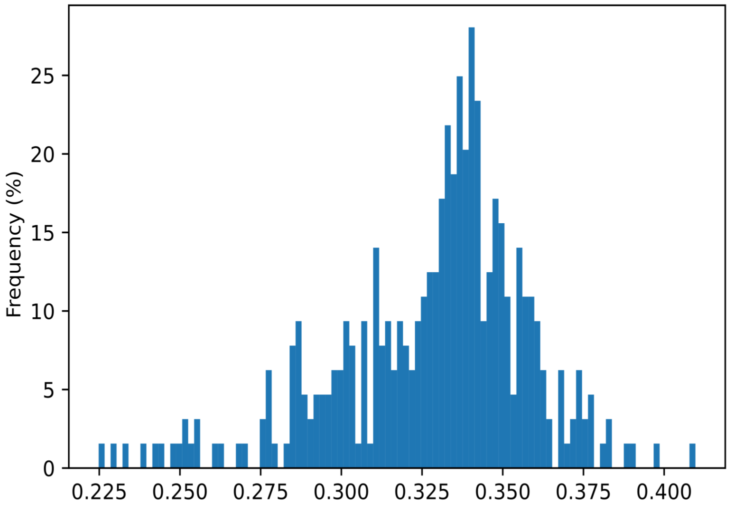


Figure S3.1: Distribution of slopes of linear regression between *lscd* and cancer incidence on a log-log scale. Of the 423 datasets from corresponding national registries in the IARC database, 347 datasets had values of cumulative incidence rate for all 17 types of cancer considered. All slopes are substantially below unity, indicating non-linear relationships; median slope = 0.334, slope range: 0.225-0.410; median intercept =-13.630, intercept range: (-15.630)-(-11.424)

Since all the data used in these papers are publicly available, we performed a model fitting exercise, taking a closer look at the nature of the associations. We considered the same 347 datasets, and found that while Pearson's r was indeed distributed around a median of 0.8 as reported by Tomasetti et al., the slopes of the regression were distributed narrowly around a median value of 0.334 (Figure S3.1). Going by the classical logic of cancer, it would mean that an average of 0.334 mutations are required to cause cancer, which is absurd. We expect a positive integer here, but get a substantially small fraction instead. This fractional slope has been identifed in passing as surprising[3], without further discussion. On the other hand, evidence in support of a non-linear relationship between cancer risk and *lscd* has been accumulating in parallel[4], but on the whole, the causal significance of such non-linearity for cancer etiology remains to be clearly elucidated. A non-linear effect of *lscd* on cancer risk either challenges the hypothesis of all required oncogenic mutations coming together purely by chance, or points to significant gaps that must be addressed.

Residuals from a regression often offer insight on the linearity of a supposed relationship, and we use this insight to investigate the purported linearity of the cancer risk-*lscd* relationship on a log-log scale. In this case, we observed that distribution of residuals around the regression was not symmetric as would be expected based on the indication of a strong linear relationship on a log-log scale. Along the first and the last one-fourth of the *lscd* range, the points lie predominantly below the line, while in the middle half of the range, a higher fraction lie above the line, as shown by the respective frequencies of positive and negative residuals (Table S3.1). In addition to the skewed distribution of points, a visual inspection of Figure S3.1 also suggested that the linear regression itself was largely driven by the first 4 points in each dataset. In line with this suspicion, upon removal of the first four points, the significance of regression was lost in all 347 datasets considered in this analysis. In order to test that the loss of significance was not due to reduction in sample size alone, the regression was performed with elimination of all other combinations of four points from each dataset ($^{17}C_4 - 1 = 2379$ combinations per dataset, not including elimination of the first four). The distribution of Pearson's r values across the combinations revealed a striking difference, with most other combinations of points retaining the significance of the relationship. This is demonstrated by a large part of the former distribution lying below the r value threshold for $p < 0.05$ ($r_{threshold} \approx 0.55$), as shown in Figure S3.2. There is therefore evidence to suggest that the log-linear relationship is largely due to the first four incidences being lower than the rest. Moreover, given the substantially less-skewed distribution of residuals about the saturation curve compared to the log-linear equation (Table S3.1), there is sufficient basis to believe that a saturation equation might in fact be a better fit to the data than a fractional power curve. Such a saturation relationship between *lscd* and cancer incidence would indicate that *lscd*, and by extension, the stem cell number, has a quantitative effect on the development of cancer only up to a certain threshold, beyond which something else becomes limiting. As we argue in the main text, these limiting factors other than the cell number and/or turnover could stem from context-dependent selection imposed through the tissue microenvironment.

Table S3.1: **Number of positive and negative residuals from the linear regression line**. The residuals were calculated from linear regressions of cumulative cancer risk, log(CR), against log(*lscd*); data were obtained from IARC as described in the text. The columns correspond to the first one-fourth, middle half and last one-fourth of the *lscd* range. Notably, a substantial skew can be seen in the extremes of the range in the linear case, with more points below the straight line than above it, reflected by a greater number of negative residuals. Compared to the linear case, the skew in the distribution of residuals is significantly lesser for the saturation equation.

| **Regression type** | **Residuals** | **First one-fourth** | **Middle half** | **Last one-fourth** |
|---|---|---|---|---|
| Linear | Positive | 667 | 1919 | 4 |
| | Negative | 1415 | 1204 | 343 |
| Saturation | Positive | 1193 | 1585 | 220 |
| | Negative | 889 | 1538 | 127 |

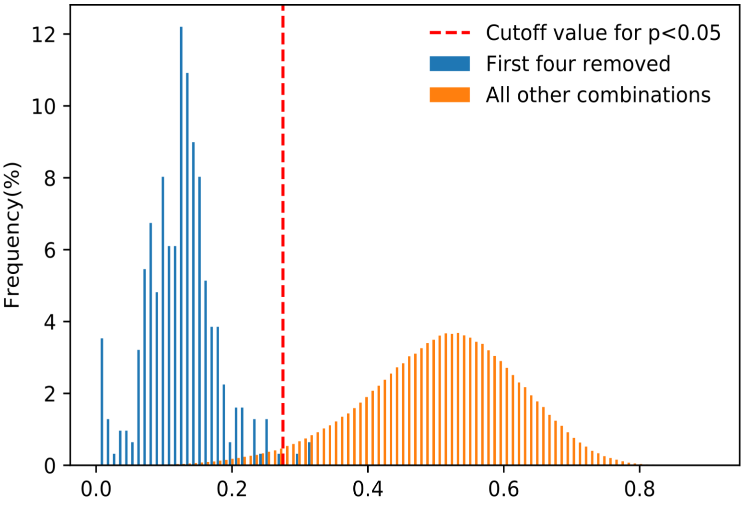


Figure S3.2: Distribution of Pearson's r for reduced datasets. The reduced datasets were obtained by removing any four data points from each sample of the IARC dataset, which have 17 data points correponding to the cancer types considered. Linear regression was performed against *lscd* for all possible combinations of removing four points from the sample, resulting in $^{17}C_4 = 2380$ regression values for every sample, each now with 13 data points. The two distributions correspond to the values of Pearon's r for the combination of the first four points removed, and those for all other combinations pooled respectively. The cutoff refers to the value of r for which $p < 0.05$, given a sample size of 13 points. The former distribution lies predominantly below this cutoff, suggesting that the linear regression is largely driven only by the first four points. This puts into question the overall inference of linearity.

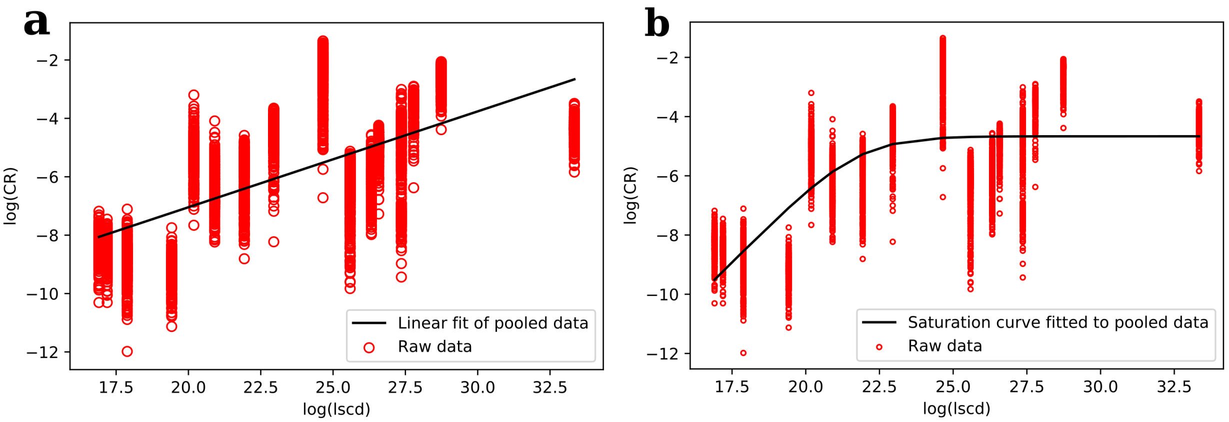


Figure S3.3: **a**. Linear vs **b**. Saturation fits. Although for statistical analyses each dataset was used separately, we show pooled data here to facilitate visual impression. Data used are the same as in previous figures.
